# Atorvastatin suppresses cardiac fibrosis and dysfunction induced by HIV and certain antiretroviral drugs in mice by blocking platelet TGFβ1

**DOI:** 10.1101/2025.09.17.676863

**Authors:** Kumar Subramani, Denys Babii, Brienne Cole, Tayyab A. Afzal, Thamizhiniyan Venkatesan, Trevor Word, Sandra Gostynska, Sixia Chen, Kar-Ming Fung, Ali Danesh, Itzayana G. Miller, Paul Klotman, Brad R. Jones, Jeffrey Laurence, Jasimuddin Ahamed

**Author notes:** To whom correspondence may be addressed. Jasimuddin Ahamed, Ph.D. **Author Contributions:** JA, JL, KS, DB, TW and BC designed the research protocol; KS, DB, TW, BC, TA, TV, SG, AD, and IGM performed the research; KS, DB, TW, KMF and SC analyzed the data; JA and JL acquired funding; TN, AN and PK, assisted in clinical conceptualization and edited the paper, JA and JL wrote the paper. **Competing Interest Statement:** The authors declare no competing interests. **Data Sharing:** For original data, please contact the corresponding Author’s lab.

## Abstract

Cardiovascular disease (CVD) both atherosclerosis-related and heart failure with preserved ejection fraction (HFpEF) and linked to cardiac fibrosis, contributes to morbidity and mortality in people with HIV (PWH) receiving antiretroviral therapy (ART). In the REPRIEVE trial, pitavastatin reduced atherosclerotic CVD risk to a magnitude inconsistent with pitavastatin’s impact solely on LDL-cholesterol and inflammation. We hypothesized that HFpEF in PWH relates to HIV-induced fibrosis mediated by platelet TGFβ1, that it is accelerated by certain contemporary ART, and may also be inhibited by statins. ART drugs used in REPRIEVE, including a nucleoside/nucleotide, integrase inhibitor-based regimen (tenofovir (TDF), emtricitabine (FTC), and dolutegravir (DTG)), and the protease inhibitors ritonavir (RTV) and darunavir (DRV), and the impact of atorvastatin, were examined in two HIV mouse models: transgenic *Tg26* mice and HIV-PDX mice engrafted with HIV-infected T cells. *Tg26* and HIV-PDX mice had higher cardiac fibrosis than littermate controls without HIV (p<0.05). Administration of TDF-FTC-DTG or RTV, but not DRV, resulted in a further ∼2-fold increase in fibrosis (p<0.01). Higher cardiac fibrosis with intracardiac fat accumulation correlated with reduced diastolic function. Mice depleted of platelet TGFβ1 (*TGFβ1*^*Platelet-Δ*^*Tg26)*, or treated with atorvastatin, were partially protected from HIV- and ART-induced cardiac fibrosis, steatosis, and diastolic dysfunction. Atorvastatin effects occurred independently of changes in inflammatory cytokines and total cholesterol. They correlated with reduced platelet activation and TGFβ1 signaling in cardiac endothelial cells, fibroblasts, and macrophages undergoing mesenchymal transition. These results indicate that certain ART regimens accelerate HIV-associated CVD characterized by HFpEF via platelet TGFβ1-dependent processes and mitigated by atorvastatin. They enhance understanding of the pleiotropic effects of statins in HIV/ART CVD and suggest a mechanism that might be targeted by antiplatelet agents or inhibition of TGFβ signaling.

**Key Points:**

- Contemporary ART regimens induce release of platelet TGFβ1 and are associated with cardiac fibrosis and diastolic dysfunction with ectopic fat deposition in HIV-infected mice.
- Depleting platelet TGFβ1 and/or treating with atorvastatin therapy suppresses HIV-ART-induced cardiac fibrosis, suggesting use of anti-platelet strategies to prevent heart failure among PWH.

## Introduction

HIV infection has been linked to several non-AIDS-defining illnesses, particularly cardiovascular disease (CVD), contributing significantly to morbidity and mortality in people with HIV (PWH) despite effective antiretroviral therapy (ART) (1). A major advance in terms of prevention derives from the Randomized Trial to Prevent Vascular Events in HIV (REPRIEVE), demonstrating that pitavastatin decreased atherosclerotic CVD among ART-treated PWH (2). The observed reduction was much greater than predicted based on the magnitude of decrease in LDL-cholesterol achieved (2). In terms of potential mechanisms, pitavastatin had no effect on other non-AIDS-defining comorbidities which, like atherosclerotic CVD, have been linked to chronic inflammation among ART-treated PWH, including end-stage liver disease, end-stage renal disease, tuberculosis, and malignancy (2, 3). Indeed, while a decrease in lipoprotein-associated phospholipase A2, a marker of arterial inflammation, was reported among REPRIEVE participants receiving pitavastatin, there was no change in other pro-inflammatory biomarkers, including hsCRP, MCP-1, sCD14, CD163, IL-1β, IL-6, IL-10, IL-18, and caspase-1 (3).

A recent secondary analysis of REPRIEVE found that pitavastatin increased procollagen C-endopeptidase enhancer 1 (PCOLCE), which enhances activity of proteinases essential for formation and assembly of vascular extracellular matrix (4). Those processes may be associated with transformation of vulnerable plaque phenotypes to more stable coronary lesions (4), consistent with the impact of pitavastatin on early atherosclerotic cardiovascular events in that trial. However, emerging data related to HIV-linked CVD in general, and the heart failure mechanisms and phenotypes that predominate among PWH in the contemporary ART era, have the potential to further modify CVD treatment and prevention strategies (5). With this concern, we sought preclinical evidence for the ability of statins to influence heart failure with preserved ejection fraction (HFpEF), which is characterized by myocardial fibrosis and steatosis in both PWH and the general population (6-10). It is now recognized in 35% of CVD cases in PWH on ART (5).

Similar to atherosclerotic CVD, the relative risk of HFpEF is increased some 2-fold among PWH treated with ART (11). In addition, while the pro-fibrotic cytokine TGFβ1 may be protective in the early stages of atherosclerosis, promoting collagen synthesis and plaque stability, the opposite has been suggested for its effects in late stages, with plasma TGFβ1 levels significantly increased in patients with coronary artery ectasia and coronary artery disease (12). Disentangling the effects of HIV vs. ART on myocardial structural and functional pathology through patient-oriented research is challenging, given potential divergent effects of different ART drugs and drug classes (13-16). In our approach, we leveraged two established mouse models of HIV: transgenic *Tg26* mice expressing seven HIV proteins but without production of infectious virus (17); and HIV patient-derived xenograft (HIV-PDX) mice, which are nonobese diabetic severe combined immunodeficiency IL2rγ^null^ mice engrafted with HIV-infected memory CD4+ T-cells from PWH (18).

We exposed these mice to ART drugs used in REPRIEVE, including a nucleoside/nucleotide and integrase strand transfer inhibitor (INSTI)-based regimen and protease inhibitors (PI) used in PI-boosted regimens. We characterized myocardial structural and functional pathology 8 weeks post-treatment. Both INSTI- and PI-based ART are associated with increased risk for many forms of CVD vs. non-nucleoside reverse transcriptase inhibitor-based ART (15, 19-22). We also tested whether atorvastatin,

lipophilic statin in the same class as pitavastatin, could protect cardiac structure/function in these settings, and explored the mechanisms of such preservation. Our data provide a novel potential pathway by which statins can influence at least certain forms of CVD in ART-treated PWH that, as in the REPRIEVE trial, occur in excess of changes in cholesterol and inflammatory cytokines.

## Results

### *HIV-Tg26* and HIV-PDX Mice Have Higher Cardiac Fibrosis Than Littermate Controls Without HIV. An INSTI-based ART Regimen and the PI, RTV Further Increased Cardiac Fibrosis

We previously showed that wild-type (wt) C57Bl/6 mice treated with supra-therapeutic doses of RTV had an approximately three-fold increase in cardiac fibrosis compared to untreated mice (∼1.5% vs. ∼0.5%) (23). Since PWH treated with PI-based ART appear at greatest risk for CVD (15, 16), including cardiac fibrosis, we first tested whether HIV alone is associated with enhanced cardiac fibrosis in mice, and whether doses of RTV equivalent to that used clinically in RTV-boosted PI regimens, or a common, INST-based ART regimen, enhance such fibrosis. Only 2 of 10 (20%) wt littermate control mice on an FVB/NJ, C57Bl/6 *or* NSG background had mild cardiac fibrosis (∼1.5% of the total cardiac area) at 4 months of age, as measured by Masson trichome and picrosirius red staining. In contrast, 8 of 11 (73%) of HIV-Tg26 mice--those containing 7 HIV-1 genes but incapable of producing infectious virus--developed mild to moderate fibrosis (2-4% of the total cardiac area). Administration of RTV or TDF-FTC-DTG, i.p. daily for eight weeks, led to moderate cardiac fibrosis (3-4% of the total cardiac area) in 12 of 15 (80%) of these mice. (P = 0.003 wt control vs. HIV-Tg26 mice and P = 0.007 in HIV-Tg26 vs. RTV-treated HIV-Tg26 mice) (**Fig.1A-B-D;** Fig. S2).

As a control, we challenged HIV-*Tg26* (**Fig. 1A-B-D**) and wt-C57Bl/6 (Fig. S2) mice with clinically relevant doses of DRV alone. This protease inhibitor has not been linked clinically to CVD in the absence of RTV boosting (15). We found no elevation of cardiac fibrosis over pre-treatment levels in the HIV transgenic or wt mice. Hematologic values (including RBC, WBC, and platelet count) showed that *Tg26* mice were mildly anemic and thrombocytopenic, but all other parameters were similar to those observed in control mice (**Table 1**). Platelet aggregation responses induced by ADP or thrombin were also similar in platelets isolated from *Tg26* and control mice (not shown).

**Table 1.** Hematologic values in control and transgenic HIV mice with treatment.

| Genotype | WBC<br>K/μL | NE<br>K/μL | LY<br>K/μL | MO<br>K/μL | EO<br>K/μL | RBC<br>K/μL | HB<br>g/dL | HCT<br>% | PLT<br>K/μL | MPV<br>fL | NE/LY<br>ratio |
| --- | --- | --- | --- | --- | --- | --- | --- | --- | --- | --- | --- |
| Control mice<br>without HIV (n=12) | 6.73±1.87 | 0.80±0.38 | 5.36±1.43 | 0.54±0.25 | 0.02±0.04 | 9.20±0.80 | 12.67±1.01 | 54.29±4.60 | 1176.67±315.80 | 4.08±0.21 | 0.15±0.07 |
| Tg26 (n=18) | 9.79±6.18 | 2.17±1.71 | 6.48±4.11 | 1.08±0.72 | 0.05±0.07 | 6.88±1.94 | 9.47±2.59 | 39.69±13.84 | 993.89±226.78 | 4.31±0.19 | 0.34±0.21 |
| Tg26 + (RTV) (n=13) | 8.08±5.59 | 2.14±2.29 | 5.14±3.08 | 0.72±0.50 | 0.03±0.06 | 8.41±1.56 | 10.12±1.73 | 46.42±8.90 | 1251.92±160.35 | 4.20±0.26 | 0.40±0.22 |
| *TGFβ1<sup>Platelet-</sup>*<br>*ΔTg26* (n=5) | 7.60±2.34 | 0.97±0.39 | 6.18±1.86 | 0.40±0.10 | 0.05±0.05 | 9.20±0.48 | 10.50±0.50 | 44.50±2.65 | 930.00±108.97 | 4.07±0.15 | 0.16±0.03 |
| Tg26 + (TDF) (n=6) | 10.97±5.03 | 3.07±2.15 | 7.02±2.92 | 0.87±0.25 | 0.02±0.03 | 7.65±1.11 | 10.50±1.95 | 43.50±6.56 | 1214.17±228.11 | 4.12±0.29 | 10.97±0.18 |
| Tg26 +<br>Atorvastatin (n=7) | 10.40±5.00 | 1.36±0.91 | 7.38±3.49 | 1.62±1.05 | 0.02±0.03 | 7.63±1.37 | 10.64±1.97 | 45.71±10.11 | 838.57±355.62 | 4.64±1.17 | 0.19±0.11 |

WBC=white blood cells; RBC=red blood cells; Hb=hemoglobin; HCT=hematocrit; PLT=platelets; MPV=mean platelet volume; NE=neutrophils; LY lymphocytes; MO=monocytes; EO=eosinophils; K=1000; M=million.

**Figure 1.**
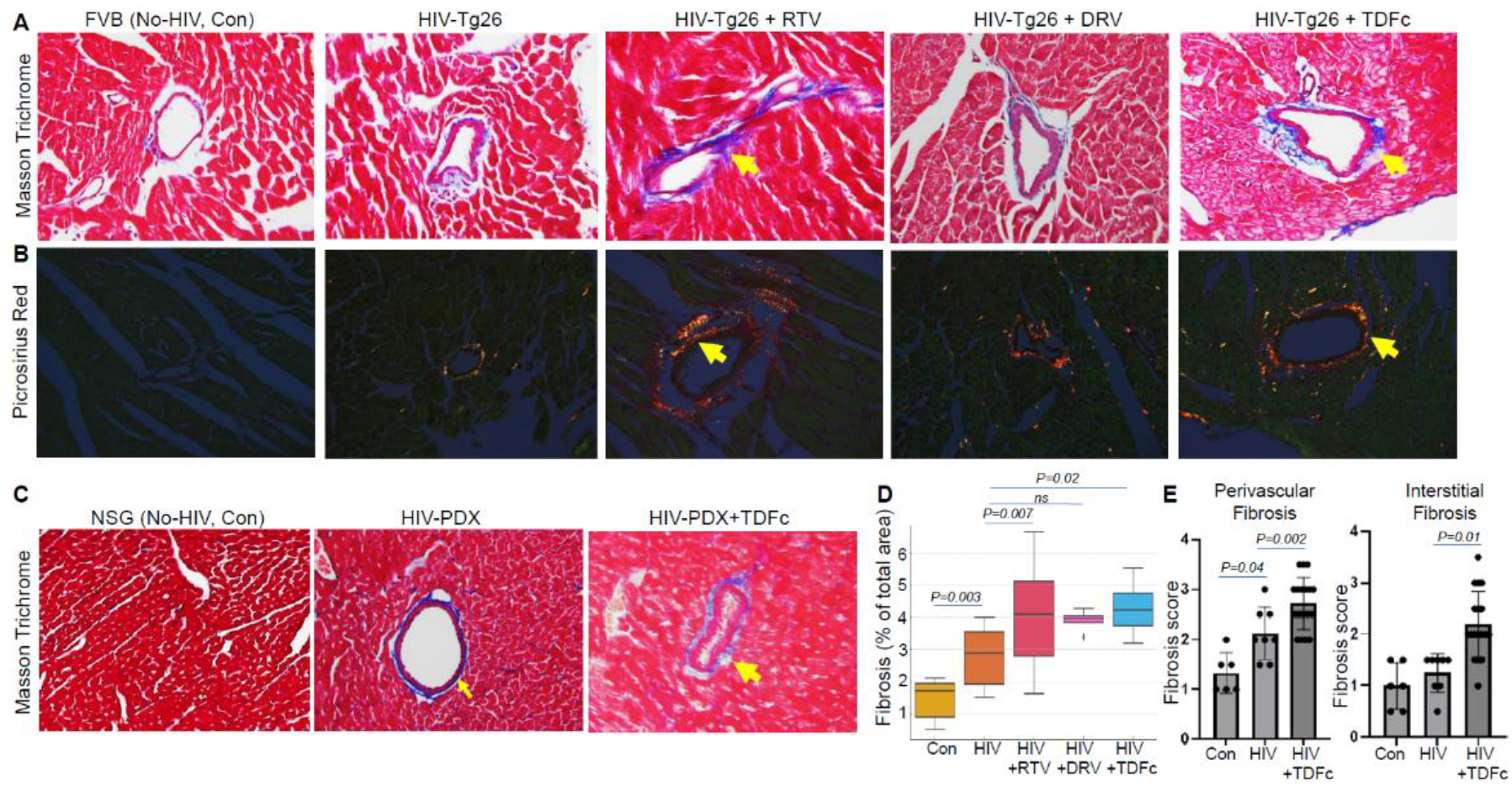
Effect of ART on cardiac fibrosis in HIV mice. (**A**) Representative images of Masson trichome staining of heart sections from FVB-No-HIV control, HIV-Tg26, and HIV-Tg26 treated with ART, ritonavir (RTV), darunavir (DRV) or with the cocktail of TDFc (TDF-FTC-DTG) for 8 weeks, showing excessive collagen in blue color and (**B**) images of picrosirius red staining, showing fibrotic areas appearing as a mixture of green, red, and yellow color, under polarized light microscope. (**C**) Representative images of Masson trichome staining of heart sections from NSG-control, HIV-infected (HIV-PDX), and HIV-PDX mice treated with the cocktail of TDFc (TDF-FTC-DTG) for 8 weeks. (**D**) Quantification of fibrotic areas from images showed that HIV mice have higher cardiac fibrosis than control mice and ART-treated mice, with the exception of DRV, had much higher fibrotic areas than those of untreated HIV-Tg26 mice. (**E**) Blind scoring for fibrosis by microscopy on a scale from 0 to 4 showed that ART-treated mice had higher perivascular and interstitial fibrosis than untreated control HIV-PDX mice.

Comparable results were observed in HIV-PDX mice infected with HIV with and without TDF-FTC-DTG treatment (**Fig.1C-E**). 9 of 11 (82%) of mice had both perivascular and interstitial fibrosis with TDF-FTC-DTG treatment vs. 20-40% of HIV-PDX mice not exposed to these drugs, as assessed by Masson trichome and picrosirius red staining and imaging by normal and-polarized light, respectively (**Fig.1C-E**).

### Both an INSTI-based ART Regimen and RTV Induce Diastolic Dysfunction with Preserved Ejection Fraction in HIV Mice

To assess whether the degree of fibrosis seen in ART-treated HIV mice was of functional significance, we assessed cardiac functions by echocardiography. Systolic dysfunction was not observed, as ejection fraction (EF) and fractional shortening (FS) were preserved in wt and combination ART-treated HIV-*Tg26* mice (Fig. S3). Diastolic function in HIV-Tg26 mice was slightly impaired over HIV negative wt controls in the absence of ART (p=0.03), but more severely impaired diastolic function was noted in both INSTI- and RTV-challenged HIV-*Tg26* vs. non-drug treated HIV-Tg26 mice, manifest by higher E/A ratio (**Fig. 2A-B,C**; p<0.01). There was a significant correlation between cardiac fibrosis and diastolic indices, specifically E/A ratios, in HIV-*Tg26* mice with and without ART vs. control *Tg26* mice (**Fig. 2D**).

**Figure 2.**
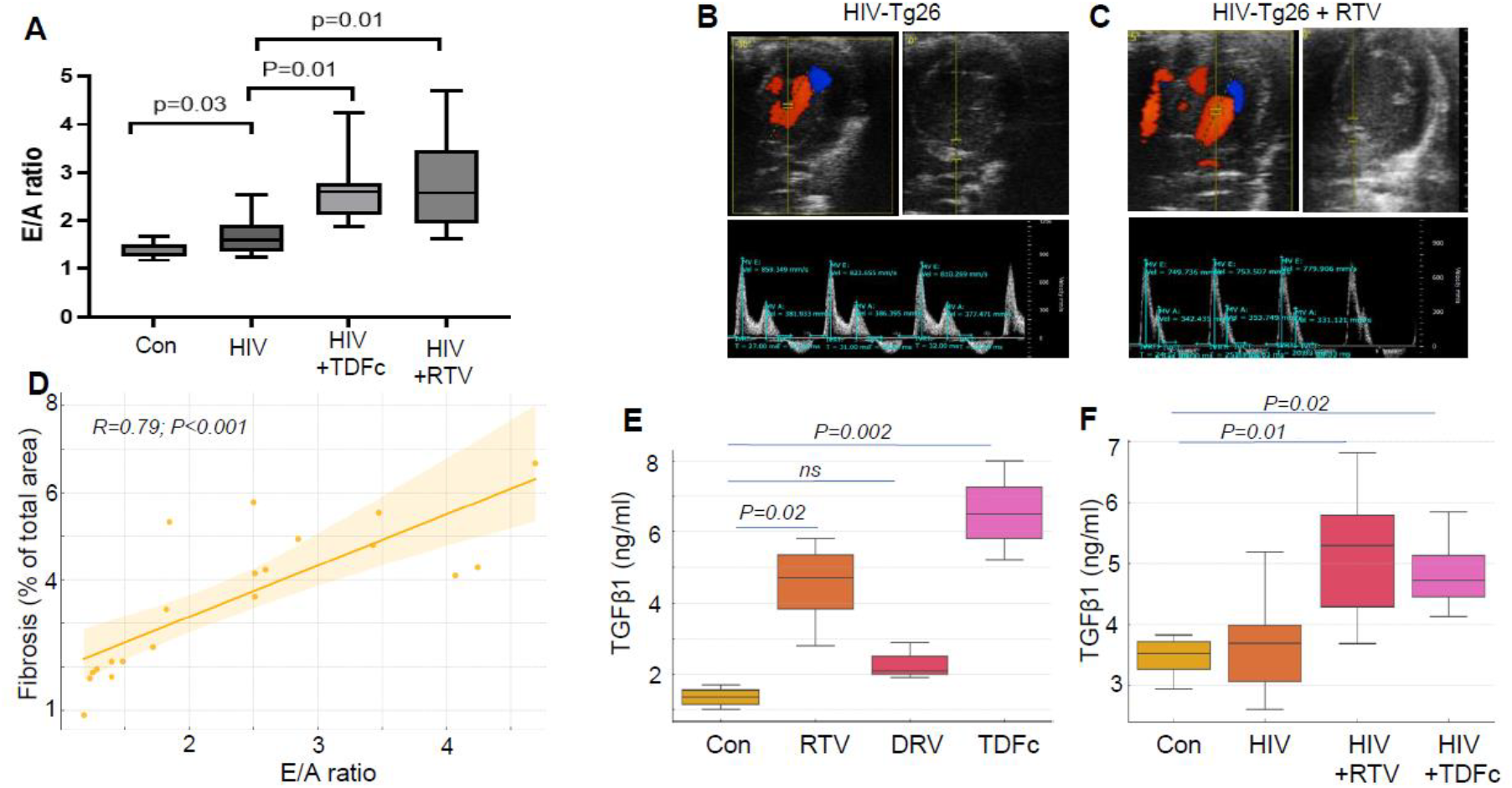
Effect of ART on cardiac function in *HIV* mice. (**A**) Cardiac function studies revealed no change in LVEF or fractional shortening following ART treatment (Fig. S3), whereas diastolic function indices, E/A ratio was higher, an indication of impairment of diastolic function, following exposure to ART, RTV or the cocktail of TDFc (TDF-FTC-DTG) for 8 weeks, vs. untreated control in HIV-*Tg26* mice. (**B-C**) Representative ultrasound images (taken using Vevo2100) of 4-chambered views with color-Doppler (upper left panels showing blood flow (red color) across the mitral valve annulus (upper panels, right) and power-Doppler (lower panel showing peak velocity of E and A waves in HIV mice without (**B**) or with RTV treatment (**C**) for 8 weeks. (**D**) E/A ratios directly correlated with cardiac fibrosis in control and ART-treated HIV mice. (**E-F**) Total TGFβ1 was measured by ELISA in platelet releasates prepared from washed platelets isolated from healthy human volunteers (10^9^per mL) before and after stimulated with RTV (5 µM) and DRV (15 µM) or with the cocktail of TDFc (TDF-FTC-DTG) at 37°C for 10 minutes and centrifuged at 14 000*g* for 10 minutes and in plasma (**F**) prepared from HIV-Tg26 mice treated with RTV or with the cocktail of TDFc (TDF-FTC-DTG).

In terms of the mechanism for this effect, we previously showed that RTV induces release of TGFβ1 in vitro from human platelets (22). We now show that RTV alone, and the TDF-FTC-DTG cocktail, induce release of TGFβ1 from freshly isolated human platelets. In contrast, the PI DRV had no effect (**Fig. 2E**). RTV and TDF-FTC-DTG also induced release of TGFβ1 in vivo, assessed in plasma from these mice (**Fig. 2F**).

### ART-Induced Ectopic Fat Deposition in the Heart is Associated with Fibrosis in HIV Mice

Fat deposition within the heart is increased in ART-treated PWH and is associated with diastolic dysfunction (19, 20). However, the relationship between fat deposition and cardiac fibrosis in the setting of HIV/ART has not been examined. We evaluated the effect of RTV and TDF-FTC-DTG on fat accumulation in the heart of HIV-*Tg26* and HIV-PDX mice. Both resulted in higher cardiac fat deposition, as measured by oil-red staining (Fig. 3A-B). To determine whether these stained areas contained fat cells, they were immunostained with an antibody to perilipin, a marker of fat cells. Clusters of perilipin-positive fat cells were identified (**Fig. 3A-B**).

**Figure 3.**
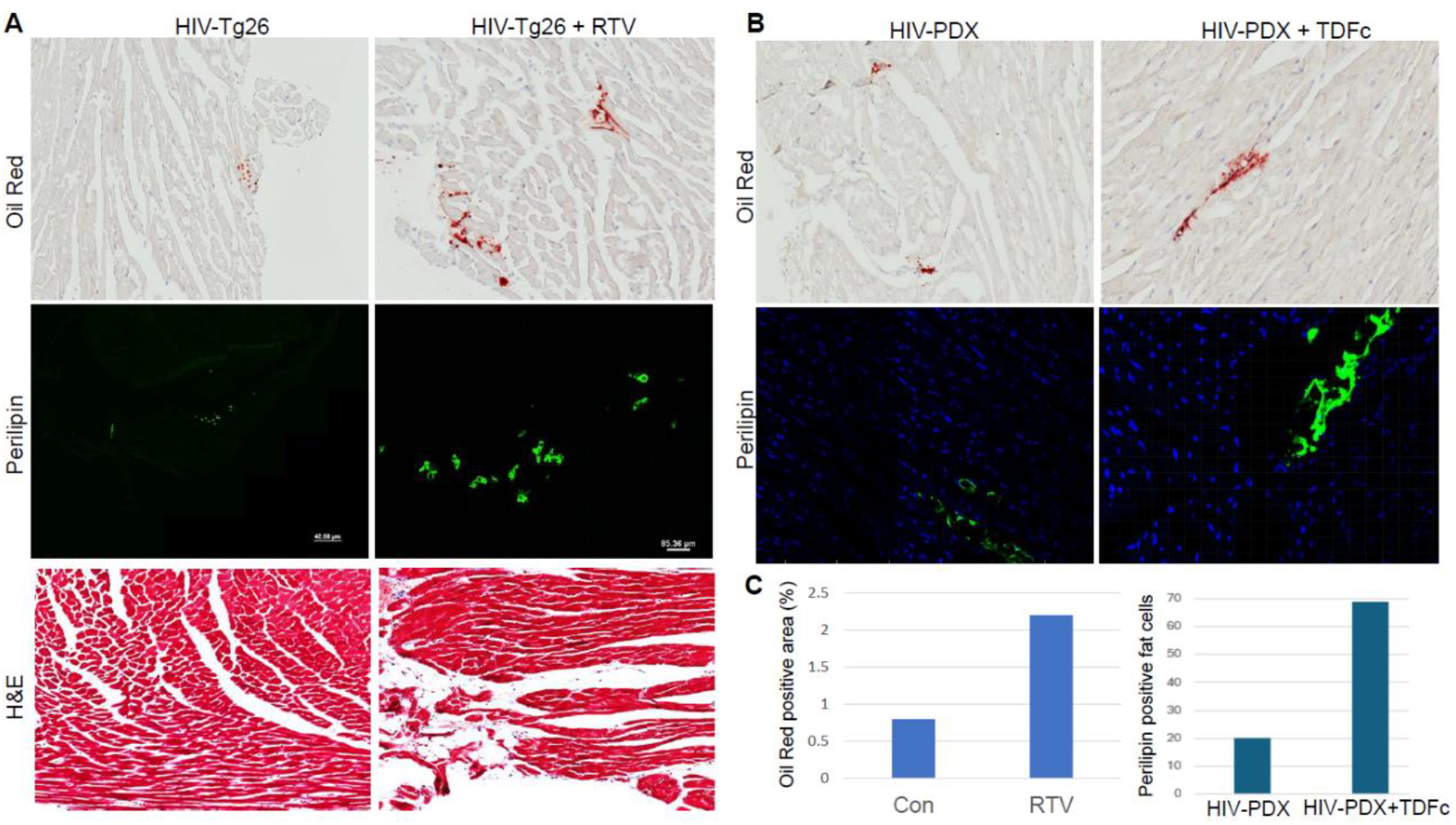
ART-induced ectopic fat deposition in the heart is associated with cardiac fibrosis in HIV mice. (**A-B**) HIV-Tg26 mice and HIV-PDX mice were challenged with either RTV alone or combination ART (TDF(c)TDF-FTC-DTG). Lipid/fat cell deposition in the heart was measured by oil-red staining (upper panels) and confirmed by immunofluorescence staining with perilipin antibody and confocal microscope imaging (middle panels). Fat cells were also confirmed by H&E staining (lower panels). Higher lipid/fat cell deposition was seen in both RTV- and combination ART-treated HIV mice. Oil-red positive areas were matched with the fibrotic areas. (**C**) Quantification of lipid/fat cells: oil-red positive area in HIV-Tg26 mice (**A**) and perilipin positive cells in ART-treated hearts of HIV-PDX mice infected with HIV (**B**).

The same heart sections were then stained with picrosirius-red. Superimposed images showed fibrotic areas with excessive collagen accumulation overlapping with fat cells. Masson trichome and H&E staining also showed fat cells surrounding fibrotic areas (**Fig. 3A**, bottom panels). Quantification revealed an association between fibrosis and lipid/fat cell accumulation in ART-treated hearts in both HIV-*Tg26* and HIV-PDX mice (**Fig. 3C**). Taken together, these results indicate that ART-treated HIV mice with higher cardiac fibrosis rates also have higher fat cell deposition in the heart.

### Platelet-derived TGFβ1 Contributes to RTV-induced Cardiac Fibrosis and Fat Cell Deposition

We previously showed that platelet TGFβ1 contributes to RTV-induced cardiac fibrosis in wt C57Bl/6 mice (23). To test whether platelet TGFβ1 contributes to cardiac fibrosis in HIV mice treated with various ART drugs, we generated *Tg26* mice with TGFβ1 deleted in platelets (*TGFβ1*^*Platelet-Δ*^*Tg26)* by crossing *Tg26* mice with *PF4CreTgfb1*^*flox/flox*^ mice. We found a >90% decrease in total TGFβ1 in platelets and serum, and a 50% decrease in plasma of *TGFβ1*^*Platelet-Δ*^*Tg26* mice compared to littermate control HIV-Tg26 mice (**Fig. 4A**). This indicates that platelets are the major source of circulating TGFβ1 in HIV mice. Although, platelets are the most abundant sources of TGFβ1, other cells, including macrophages, dendritic cells, regulatory T cells, and B cells can also produce TGFβ1 (24).

**Figure 4.**
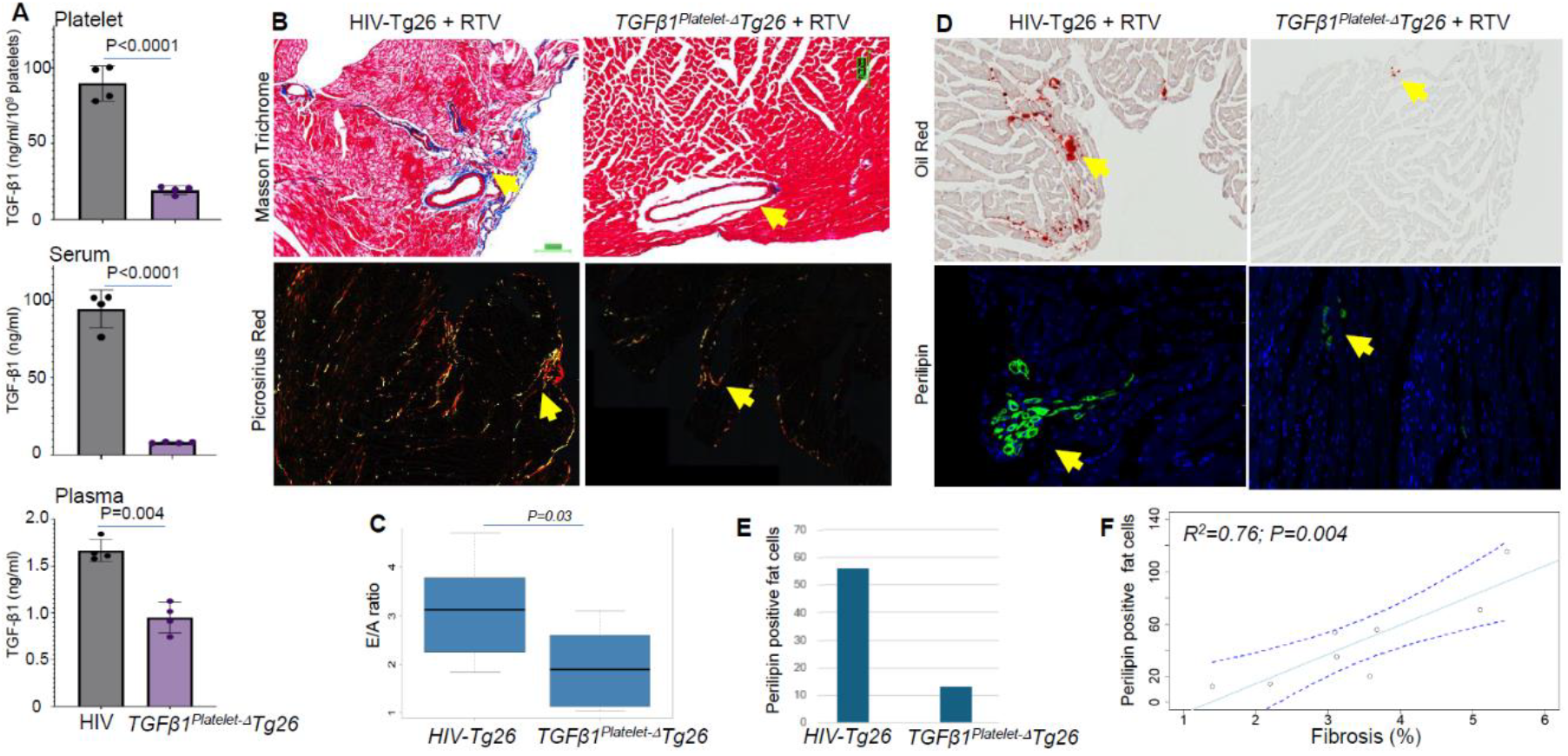
*TGFβ1*^*Platelet-Δ*^*Tg26* mice are partially protected from developing RTV-induced cardiac fibrosis and deterioration of diastolic function. (**A**) *TGFβ1*^*Platelet-Δ*^*Tg26* mice had >90% less TGFβ1 in their platelets and serum and >50% less in plasma than their littermate control HIV-Tg26 mice, as measured by ELISA (n = 4). (**B**) Masson trichrome and picrosirius red staining of heart sections of RTV-challenged HIV-*Tg26* or *TGFβ1*^*Platelet-Δ*^*Tg26* mice. Fibrotic areas in blue (Masson trichrome images) and a mixture of green, red, and yellow, fluorescent color (picrosirius red images taken under polarized light) showed lower collagen accumulation in *TGFβ1*^*Platelet-Δ*^*Tg26* than HIV-Tg26 mice. Quantification of fibrotic areas from images taken with a polarized light microscope showed that *TGFβ1*^*Platelet-Δ*^*Tg26* mice had smaller fibrotic areas than those of HIV-*Tg26* littermate control mice (percentage of picrosirius red–stained fibrosis areas were 2.5% ± 0.5% in *TGFβ1*^*Platelet-Δ*^*Tg26*, 3.3% ± 0.7% in Tg26 mice; *P* <0.01). (**C**) Diastolic heart functions (E/A ratio) were measured by echocardiography, showing protection of impairment from diastolic dysfunction in *TGFβ1*^*Platelet-Δ*^*Tg26* compared to HIV-*Tg26* mice. (**D**) Lipid/Fat cells accumulation in heart tissue was measured by oil-red and perilipin staining, showing lower fat cells accumulation in *TGFβ1*^*Platelet-Δ*^*Tg26* mice than in HIV-*Tg26* after RTV challenge. (**E**) Number of perilipin-positive fat cells (green color) per entire heart, with images taken by confocal microscope. (Each bar represents the average value from five heart sections). (**F**) Correlation between fibrosis and fat cell deposition in ART-treated hearts of *TGFβ1*^*Platelet-Δ*^*Tg26 and HIV-Tg26* mice challenged with RTV.

Higher amounts of cardiac fibrosis were observed in RTV-treated *Tg26* littermate mice vs. RTV-treated *TGFβ1*^*Platelet-Δ*^*Tg26* mice (**Fig. 4B**). The percentage of fibrosis areas was 2.5% ± 0.5% in *TGFβ1*^*Platelet-Δ*^*Tg26* mice vs. 3.3% ± 0.7% in HIV-Tg26 mice; P <0.01). This difference in fibrosis was paralleled by protection from impairment of diastolic function, as seen in *TGFβ1*^*Platelet-Δ*^*Tg26* mice treated with RTV (**Fig. 4C**). Since ART-induced accumulation of lipid/fat cells in the heart correlated with fibrosis, we also assessed lipid/fat cell accumulation in the *TGFβ1*^*Platelet-Δ*^*Tg26* mice. Staining of RTV-treated *TGFβ1*^*Platelet-Δ*^*Tg26* mouse hearts with oil-red revealed lower lipid accumulation compared to HIV-*Tg26* littermates challenged with RTV. Immunostaining with perilipin confirmed less accumulation of fat cells in these mice compared to littermate controls (**Fig. 4D, E**). The presence of fat cells correlated with levels of fibrosis in *TGFβ1*^*Platelet-Δ*^*Tg26* and Tg26 littermate control mice challenged with RTV (**Fig. 4F**). To test whether ART-associated release of TGFβ1 *in vivo* is primarily derived from platelets, we challenged *TGFβ1*^*Platelet-Δ*^*Tg26* mice with RTV and TDF-FTC-DTG. No increase in TGFβ1 levels over baseline was observed (data not shown), indicating that ART increases TGFβ1 in vivo primarily by activating platelets.

### Atorvastatin Suppresses TGFβ Signaling, Cardiac Fibrosis and Fat Accumulation, and Diastolic Dysfunction Without Altering Inflammatory Cytokines and Lipid in HIV-Transgenic Mice

The REPRIEVE trial showed that pitavastatin, a lipophilic statin, had a beneficial effect in reducing atherosclerotic CVD (2). We first tested whether a related lipophilic statin, atorvastatin, could influence processes related to the second major type of CVD linked clinically to HIV/ART, fibrosis-associated HFpEF, by inhibiting TGFβ1-induced signaling. We used an engineered cell culture system in which TGFβ1 stimulation induces PAI1 luciferase activity in mink lung epithelial cells (MLEC) expressing a TGFβ-responsive PAI1 reporter fused with a luciferase gene (25). This signaling response was inhibited by atorvastatin in a dose-dependent manner (**Fig. 5A**). This is consistent with the finding that TGFβ1-induced SMAD2/3 phosphorylation signaling was inhibited by atorvastatin, also in a dose-dependent manner (**Fig. 5B-C**). Atorvastatin also inhibited TGFβ1-induced signaling responses for PAI1 and SMAD phosphorylation in vitro in the presence of RTV or TDF (Fig. S5A-B-C).

**Figure 5.**
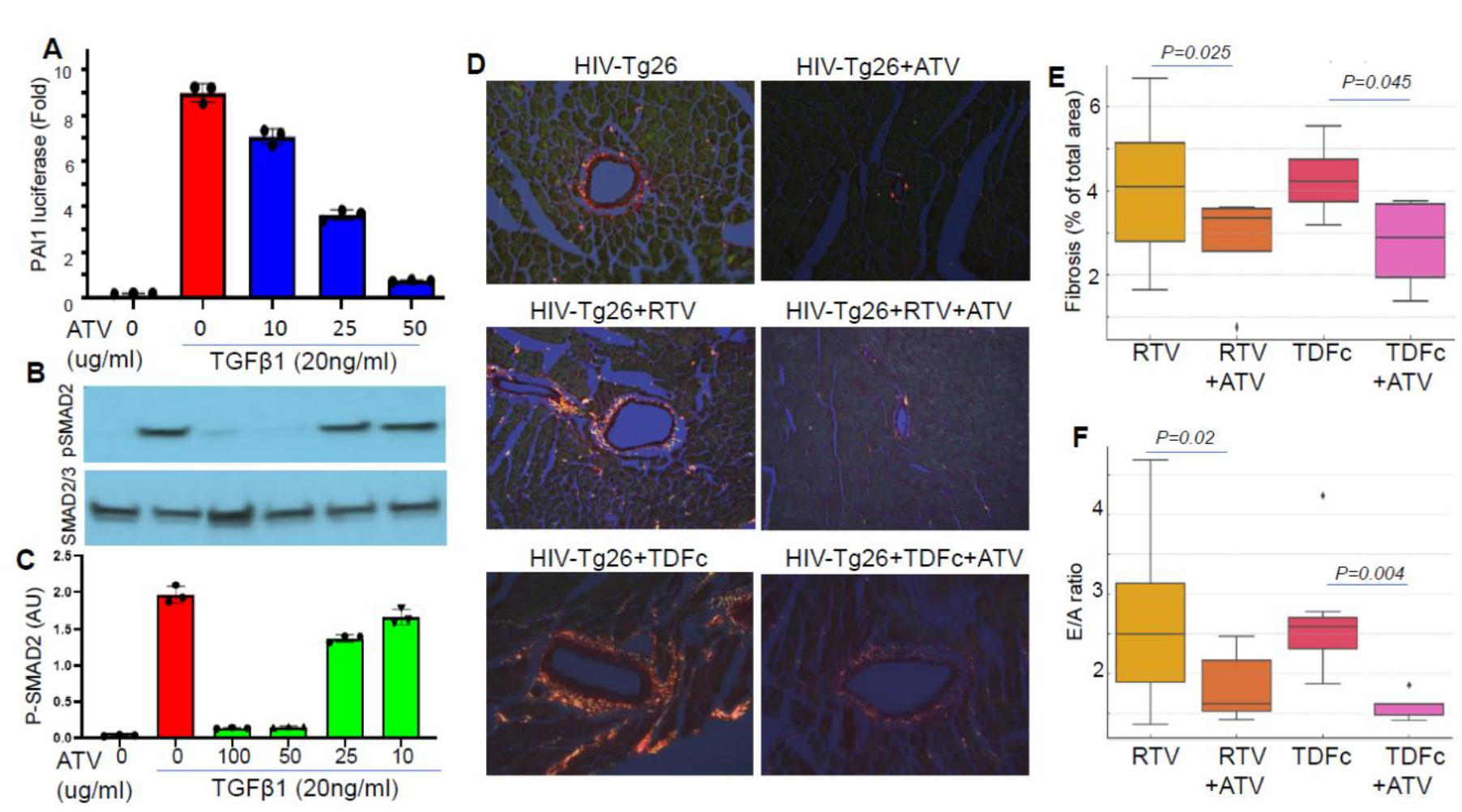
Atorvastatin (ATV) suppresses TGFβ1 signaling, cardiac fibrosis, and fat cell deposition and improves diastolic dysfunction in HIV mice. (**A**) ATV inhibited TGFβ1-mediated signaling for PAI-1 luciferase activity and (**B**) SMAD2 phosphorylation in a dose-dependent manner in mink lung epithelial cells stimulated with platelet TGFβ1 (20ng/ml) for 16-18h. (**C**) Quantification of SMAD2 phosphorylation. (**D**) Picrosirius red staining images (taken by polarized microscope) of heart sections of HIV-Tg26 mice challenged with vehicle, or RTV, or the cocktail of TDFc (TDF-FTC-DTG) and with ATV for 8 weeks, showing ATV halted cardiac fibrosis. (**E**) Quantification of fibrotic areas showed lower fibrosis in ATV than vehicle co-treated RTV- or TDFc-exposed mice. Comparison of among groups using Wilcoxon rank-sum tests, the ATV co-treated significantly reduced fibrosis levels. (**F**) Diastolic function (E/A ratio) was measured by echocardiography, showing less impairment in ATV-vs. HIV-*Tg26* mice exposed to RTV or TDFc.

To investigate whether atorvastatin could prevent HIV-linked cardiac fibrosis in HIV mice in vivo, we administered 3 mg/kg in drinking water to HIV-*Tg26* mice and measured cardiac fibrosis and heart function. This dose corresponds to 20 mg/day in an 80-kg human and is well-tolerated in mice, without toxicity (26, 27). Less cardiac fibrosis and impairment of diastolic function was observed compared to untreated HIV-*Tg26* mice over an eight-week period (**Fig-5D-E**). Atorvastatin administration along with TDF-FTC-DTG or RTV to *Tg26* mice also reduced cardiac fibrosis and preserved diastolic function compared to ART-exposed *Tg26* mice not receiving atorvastatin (**Fig. 5D-F;** Fig. S6). Atorvastatin treatment does not alter inflammatory cytokines IL-6 and TNFα or plasma total cholesterol and glucose levels in HIV-Tg26 mice with or without ART treatment (Fig. S4). Staining of heart sections with Masson trichome, Picrosirius-red, or Oil-red and perilipin revealed lower fibrosis and lipid/fat cell accumulation in atorvastatin-treated vs. vehicle treated mice challenged with TDF-FTC-DTG or RTV (Fig. S6).

### Atorvastatin Inhibits Platelet Activation and TGFβ Signaling and Suppresses Myofibroblast Transition and Collagen Expression in Cardiac Cells

It has been shown that atorvastatin inhibits platelet activation induced by classic platelet agonists (28). We now show that this drug inhibited human platelet activation in vitro as well as reduced plasma TGFβ1 levels in mice associated with certain antiretroviral drugs (Fig. S7).

To test whether atorvastatin also blocks TGFβ signaling in vivo, we performed multicolor immunofluorescent staining and confocal imaging of HIV mouse hearts with and without atorvastatin treatment. HIV-*Tg26* mice had higher TGFβ signaling, documented by the increased phosphorylation of SMAD2 and SMAD3 (pSMAD2/3), in cells within areas of perivascular fibrosis (**Fig. 6** and Fig. S8). Cells positive for pSMAD3 were also positive for αSMA and/or collagen, or periostin (**Fig. 6A**), indicating that they were undergoing mesenchymal transition to myofibroblasts, cells known to produce excessive collagen in tissue undergoing pathologic fibrosis (29). HIV*-Tg26* mice treated with atorvastatin had significantly lower levels of nuclear pSMAD2/3 translocation in cells co-expressing myofibroblast markers (**Fig. 6** and Fig. S8), suggesting that atorvastatin inhibits TGFβ1-induced signaling for fibrosis-related gene responses *in vivo*.

**Figure 6.**
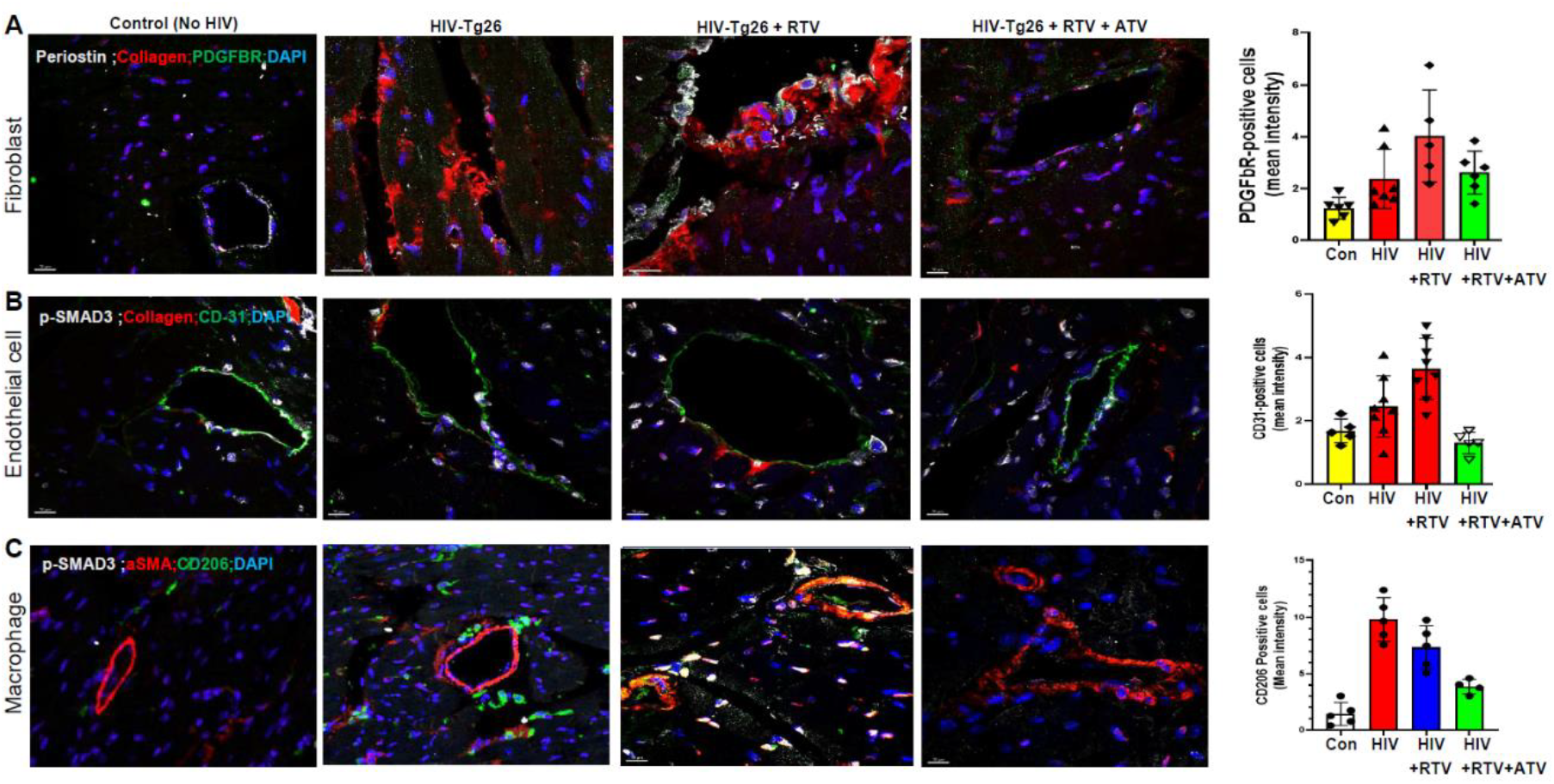
Atorvastatin inhibits Smad signaling and mesenchymal transition of fibroblasts, endothelial cells, and macrophages into myofibroblasts in heart of HIV mice. Confocal images and quantification of (**A**) PDGFβR-positive fibroblasts (**B**) CD31-positive endothelial cells, and (**C**) CD206-positive macrophages in hearts of HIV-*Tg26* mice expressing higher αSMA or collagen vs. controls (No-HIV), with or without RTV exposure. Atorvastatin (ATV) suppressed these effects. Fig. S6 shows images and quantification of TGFβ signaling response assessed by pSmad3, along with CD31.

To further identify cells contributing to mesenchymal transition, producing excessive collagen via inducing TGFβ1 signaling and cardiac fibrosis, we co-stained heart sections with fibroblast (PDGFβR)-(**Fig. 6A**), endothelial cell (CD31)-(**Fig. 6B**), and macrophage (CD206)-(**Fig. 6C**) specific markers and antibodies to α-SMA and/or collagen or periostin. We observed higher numbers of cells expressing CD31-, PDGFbR-, and CD206-, and collagen or α-SMA or periostin along with pSMAD3 in HIV-Tg26 mice treated with RTV or TDF-FTC-DTG, whereas fewer cells with colocalization of these markers were observed in the atorvastatin-treated hearts (**Fig. 6A-B-C**). Quantification of collagen–positive areas showed lower levels in ATV-treated hearts compared with their vehicle-treated HIV-Tg26 mice (**Fig. 6**), indicating these cells are transitioning into myofibroblasts and atorvastatin reduced these effects. In addition, TGFβ1 induced *ACTA2* (alpha-smooth muscle actin) and *COL1A1* (collagen) gene expression, markers of myofibroblasts, which was also blocked by atorvastatin (Fig. S5D).

Taken together, these data indicate that platelet-derived TGFβ1-mediated signaling in cardiac cells (endothelial cells, fibroblasts, macrophages, or other unidentified cells) can trigger their transformation into collagen-producing myofibroblasts, inducing cardiac fibrosis. These processes can be mitigated by atorvastatin or by inhibition or depletion of platelet TGFβ1, as illustrated in **Fig. 7**.

**Figure 7.**
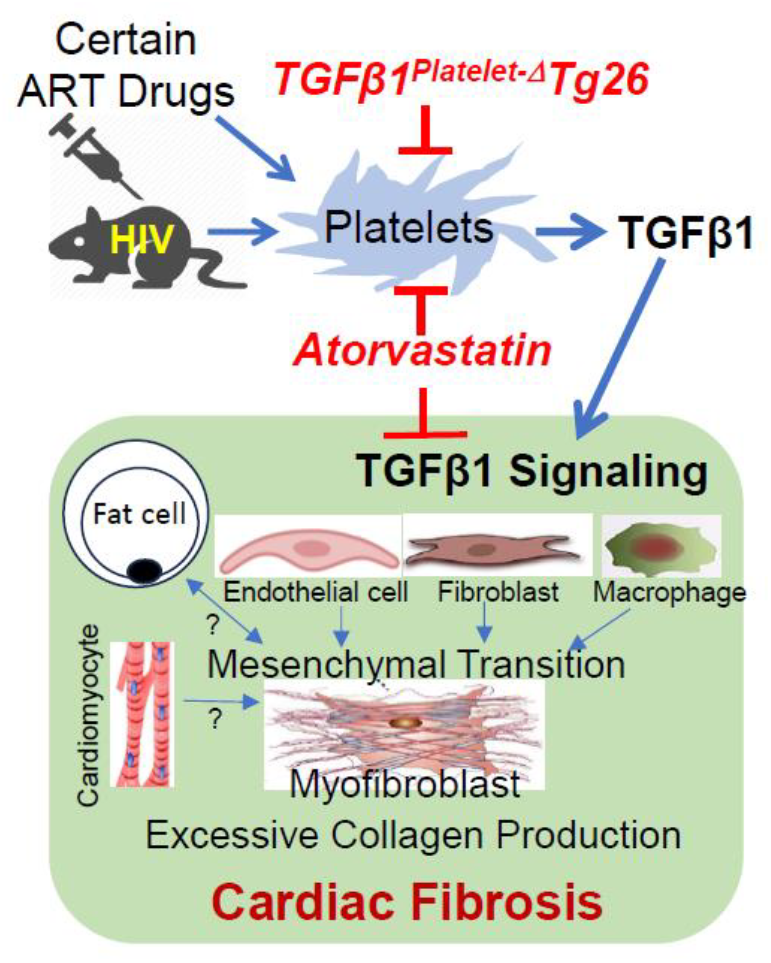
Graphic model showing that HIV-ART induces release of TGFβ1 from platelets, inducing TGFβ signaling in different cardiac cells, undergoing mesenchymal transition into myofibroblast, producing excessive collage, leading to cardiac fibrosis. Mitigation of these responses by atorvastatin inhibiting TGFβ signaling and platelet activation.

## Discussion

Our work investigating HIV/ART-induced cardiac pathology in mouse models of HIV infection yielded several novel findings of clinical relevance to accelerated HFpEF seen in PWH. We had previously shown that platelet-derived TGFβ1 is an important mediator of RTV-associated cardiac fibrosis in wt mice (23). We now document that a lower dose of RTV, as used in PI-boosted ART regimens, but not the PI DRV alone, and the INSTI-based regimen TDF-FTC-DTG, augment cardiac fibrosis and impaired diastolic function in the context of HIV, above levels seen with HIV alone. The level of cardiac fibrosis seen in HIV/ART mice, 3-4%, was equivalent to that estimated by imaging studies in ART-treated PWH (2-3%) (30). It was sufficient to induce diastolic dysfunction. It was accompanied by cardiac steatosis, also a feature of HFpEF in ART-treated PWH. Areas of ART-induced fibrosis were accompanied by myofibroblast infiltration with elevated levels of SMAD2/3 phosphorylation, reflecting TGFβ1 signaling and excess collagen production. A role for platelet TGFβ1 in cardiac fibrosis associated with certain ART drugs was further supported by the correlation of plasma TGFβ1 levels with both fibrosis and diastolic function.

In terms of practical interventions, atorvastatin protected *Tg26* mice, both untreated and exposed to either RTV or the INSTI-based cocktail TDF-FTC-DTG, from cardiac fibrosis and steatosis and diastolic dysfunction. These effects appear to be channeled, at least in part, through inhibition of TGFβ signaling. They correlated with the presence of fewer myofibroblasts in the hearts of HIV/ART mice treated with atorvastatin vs. vehicle. These results in atorvastatin-treated HIV/ART mice are consistent with the known antioxidant/anti-inflammatory effects of statins in general (31), and the fact that atorvastatin can attenuate the rise in soluble ST2, a marker of fibrosis, in PWH on ART (32). Although pitavastatin has a greater effect on HDL-cholesterol than atorvastatin (33), the latter can significantly reduce non-calcified coronary plaque volume relative to placebo in PWH on ART, despite no change in HDL-cholesterol (34). It also has an indirect immunomodulatory effect in the setting of HIV, increasing Treg and inhibiting expression of the critical HIV coreceptor CCR5 (35). Finally, although potential drug-drug interactions with statins and ART are primarily based on theoretical considerations, caution is recommended using atorvastatin, but not pitavastatin, in PI-based ART due to effects on cytochrome CYP3A4 (36). However, NRTIs and INSTIs share an affinity to the BCRP (breast cancer resistance protein) drug transporter with pitavastatin, while no such interaction occurs with atorvastatin (36). Selection of a specific statin in CVD prophylaxis for PWH on ART might be individualized based on these and other more traditional CVD risk considerations.

Our observations dovetail with recent work by Marunouchi et al. demonstrating that statins suppressed cardiac fibrosis and diastolic dysfunction in wt mice fed a high-fat diet and exposed to the nitric oxide synthetase inhibitor N[w]-nitro-L-arginine methyl ester hydrochloride as a chemical means of inducing HFpEF (37). Assessment of cardiomyocytes from statin pre-treated vs. control mice revealed decreased phosphorylation of SMAD and MAPK, enzymes downstream of TGFβ, reflecting reduced TGFβ signaling in those cells (37). Our in vitro data showing that atorvastatin dose-dependently inhibits TGFβ1-induced SMAD2/3 signaling and TGFβ-responsive profibrotic responses, including PAI1 and αSMA, are consistent with studies in human heart fibroblasts showing that atorvastatin suppresses SMAD and MAPK signaling (38). Our data also extend the work of previous studies (28) indicating that atorvastatin can block platelet activation linked to standard agonists (ADP, collagen, arachidonate) and certain cytokines. It is thus plausible that stains work via a dual mechanism in HIV/ART-associated CVD characterized by HFpEF. blocking TGFβ1 release from platelets and subsequently TGFβ signaling in cardiac cells.

In terms of the impact of atorvastatin on cholesterol and inflammation, traditional CVD risk factors, statins can lower inflammatory cytokines and LDL-cholesterol in humans, but those effects did not correlate with the ability of a related lipophilic statin, pitavastatin, to impact atherosclerotic CVD in ART-treated PWH (2). Those results are paralleled by our mouse models, as no significant reduction in IL-6, TNF-α, or cholesterol was seen. We acknowledge that our data does not provide conclusive evidence that atorvastatin’s effects were completely independent of modulation of cholesterol metabolism.

Our focus on TGF-β and fibrosis should also be viewed in the context of HIV/ART CVD risk in general, both atherosclerotic and fibrosis-associated. In REPRIEVE, pitavastatin increased abundance of PCOLCE, which enhances proteinases involved in vascular extracellular matrix production, favoring transformation of plaques vulnerable to fragmentation to more stable coronary lesions (4). This would be consistent with the salutary effects of TGF-β1 in the early stages of atherosclerosis (12). In later stages, with development of pathologic fibrosis and HFpEF, suppression of TGF-β1 may be beneficial. Both stages would then benefit from statin intervention, but for distinct reasons. Although there are no validated tools to predict who among PWH receiving ART may clinically progress to HFpEF, the newly developed American Heart Association PREVENT HF risk score has proven of value in a limited clinical study of such individuals(11).

The concomitant development of cardiac steatosis with fibrosis in our ART-treated HIV mice is also of interest. Among PWH in REPRIEVE, increased peri-coronary adipose tissue density was independently associated with prevalence and severity of coronary plaque (39). But the interplay between cardiac fibrotic and steatosis pathways is incompletely understood. While ART-treated PWH without heart failure have higher levels of cardiac fibrosis and steatosis vs. individuals without HIV, cardiac steatosis stands out as the structural pathology correlating most closely with diastolic dysfunction (9, 30, 40, 41). Cardiac magnetic resonance spectroscopy-based studies reveal excess intramyocardial lipid deposition among ART-treated PWH in association with reduced diastolic function (9, 40). Certain ART drugs are also associated with ectopic visceral and hepatic fat deposition (42). Our findings underscore the need for further exploration of HIV/ART-induced cardiac fat accumulation. Cardiac steatosis is an emerging problem among ART-treated PWH globally (1, 20), paralleling rising rates of obesity in this population (43). Future studies might assess whether anti-obesity therapeutics known to confer cardioprotection, such as glucagon-like peptide 1 receptor agonists (GLP-1 RA) (44), reduce cardiac and other ectopic fat accumulation among PWH. In this regard, a pilot study in which ART-treated PWH received the GLP-1 RA semaglutide revealed weight loss accompanied by a significant reduction in liver fat (45).

Our study has limitations. Based on our prior work with RTV, we focused our mechanistic experiments on TGFβ pathways leading to fibrosis. However, other pathways may also be relevant. Our work was strengthened by study of ART-induced cardiac pathology in two different mouse models of HIV, as well interrogation of ART effects among mice with platelet-specific TGFβ1 deletion to gain mechanistic insights. We also explored the impact of concomitant statin therapy, paving the way for clinical correlations. Selection among classes of ART drugs, and individual drugs within classes, based on data presented here, could be considered. Apart from statins, suppression of ART-induced TGFβ1 release from platelets and its activation could be explored. This might be accomplished via anti-platelet agents, compounds that inhibit TGFβ1 released from platelets, and pharmacological inhibition of TGFβ signaling, such as with galunisertib (46).

## Materials and Methods

### Mouse Experiments

Mouse experiments were conducted in accordance with the NIH Guide for the Care and Use of Laboratory Animals and were approved in advance by the Institutional Animal Care and Use Committee. Each mouse was numbered, and all experiments were performed by investigators blinded to the genotype of the mice. All mice were housed in a controlled environment (23°C ± 2°C; 12-hour light/dark cycles) and fed a standard diet (PicoLab Rodent Diet). We studied both male and female mice, aged 6– 10 weeks. *HIV-Tg26* is a transgenic mouse that expresses seven of the nine HIV proteins under the control of a viral LTR promoter(17). They were bred with either FVB/NJ mice or C57Bl/6 background for at least 10 generations. To create a platelet-specific knockout of the *Tgfb1 gene* on a *Tg26 background*, we crossed PF*4CreTgfb1*^*flox/flox*^ *(TGFβ1*^*Platelet-Δ*^) mice(29) *with Tg26 mice to generate TGFβ1*^*Platelet-Δ*^*Tg26* mice. *We confirmed inactivation of th*e *Tgfb1*-floxed allele in platelets by genotyping and performed phenotypic characterization by measuring TGFβ1 levels in platelets, serum, and plasma in mice homozygous for *TGFβ1*^*Platelet-Δ*^*Tg26. Tg26* littermates without *PF4Cre* or *Tgfb1*^*flox*^*/*^*flox*^ were used as controls.

Humanized HIV-PDX mice were generated by engraftment of memory CD4+ T cells to NSG mice (NOD/SCID IL2rγ^−^/^−^, Jax #00557). Memory CD4+ T cells were isolated from PBMCs of a person living with HIV on cART(18). Mice received memory CD8+T cells from the same donor on day 35 post-engraftment, and were then inoculated i.v. with 10,000 TCID50 of the JRCSF isolate of HIV-1. The mice were treated subcutaneously from day 84 with 100uL of vehicle or a combination ART regimen consisting of TDF (57mg/kg), FTC (143mg/kg), and DTG (7mg/kg)(47), for 4-8 weeks. All HIV-infected mice were housed in the Belfer Research Building animal facility of Weill Cornell Medicine.

### Antiretroviral Drug Dosing

Dosages of ART drugs (RTV 5.5mg/kg, DRV 16.2mg/kg) used parallel those used in humans, as documented by PK/PD data in mice(48) and measurements of antiretroviral drug concentrations in lymph nodes of humanized mice vs. humans, showing equivalent tissue levels(49). Of note, our ART medications involved much lower concentrations than used in many published rodent models of HIV infection (50), where effects may have been influenced by supra-therapeutic dosing.

### Collection and Preparation of Mouse Serum, Plasma, and Platelet Releasate

To obtain serum, whole blood was collected by retrobulbar puncture (RB) in tubes without an anticoagulant and incubated at 37°C for 4 hours. Serum was collected as the supernatant after centrifuging samples at 13,000*g* for 20 minutes at 4°C. To obtain plasma, blood was drawn by retrobulbar (RB) puncture and placed in a polypropylene tube containing 0.1 volume of 3.8% sodium citrate and PGE1 (1μM), pH 7.4. Plasma was isolated by centrifuging samples immediately after blood drawing at 12,000*g* for 5 minutes at 4°C. All samples were stored at -80°C until analysis. Washed platelets were prepared as we previously described(51). Platelets (0.5 × 10^9^) were stimulated with drug for 10 minutes at 37°C and releasates collected as the supernatants following sample centrifugation at 13,000*g* for 15 minutes at 4°C.

### Measurement of TGFβ1 Levels and Signaling

Total TGFβ1 in platelets, serum, and plasma was measured after converting latent TGFβ1 to active TGFβ1 by acidification followed by neutralization, using a DUO-antibody ELISA assay specific for the activated form of TGFβ1 (R&D Systems). TGFβ1-induced signaling activity was measured by a functional bioassay using a mink lung epithelial cell (MLEC) line stably expressing a luciferase reporter gene under control of the plasminogen activator inhibitor 1 (PAI1) promoter(25). Briefly, MLECs (2.5 × 10^4^) were plated in a 96-well tissue culture plate and allowed to adhere for 3 hours. The medium was replaced with 90 μL serum-free medium (DMEM containing antibiotics), and the sample to be tested (10 μL) added and incubated for 16-18 hours at 37°C. Luciferase activity driven by the PAI1 promoter was assayed from cell lysates in an automated luminometer using a luciferase assay system (Promega, Madison, WI). The MLEC assay was used to confirm that active TGFβ1 was driving the signaling response. We also assessed TGFβ1 signaling via Smad activation by stimulating mouse endothelial cells or MLEC with active TGFβ1 and immunoblotting for Smad2/3 phosphorylation using mAb specific for phosphorylated Smad2/3. In some experiments, samples were incubated with a TGF-β1 neutralizing antibody or atorvastatin to assess the specificity of TGFβ1 signaling detected by the PAI1 luciferase and Smad2 phosphorylation assays.

### Assessment of Cardiac Function

Systolic and diastolic functions were measured with a high-resolution ultrasound system Vevo 2100® (VisualSonics) using established methods(51). Systolic function parameters were assessed at end-diastole (d) and end-systole (s) using M-mode images at the level of the papillary muscles in a left parasternal short axis view. LV ejection fraction (EF) and fractional shortening (FS) were calculated. Diastolic function indices were recorded in an apical 4-chamber view sound window using pulse-wave or tissue Doppler echocardiography to measure E- and A-wave peak velocities. A’- and e”-tissue Doppler waves and E/A ratio was calculated following the method described previously(51).

### Histology

Animals were euthanized, and the hearts were excised, perfused with saline, weighed, and fixed in 4% paraformaldehyde. Myocardial fibrosis was evaluated by staining with hematoxylin and eosin, Masson trichrome, and picrosirius red. Grading of fibrosis on a scale of 0 to 4 was performed by an expert veterinary pathologist without knowledge of the treatment using methods established in our lab(51). Images were taken through high magnification assessment of Masson’s trichrome-stained images taken by Aperio Slide Scanner, and picrosirius red staining images were taken in a polarized scanner. An artificial intelligence-deep learning method was adapted to set an algorithm for quantification(52) (Fig. S1).

### Oil-red Staining for Cardiac Fat/Lipid

Heart sections were fixed with 4% paraformaldehyde for 24h followed by OCT mounting and cryo-sectioning (6µ thickness). They were stained with oil-red stain (Millipore), following the manufacturer’s protocol. Pictures were taken by scanning whole heart sections using an Aperio Slide Scanner and assembled for quantification using Image scope software.

### Immunostaining and Confocal Imaging

Immunofluorescence-stained OCT or parrafin-embeded heart sections (5-6µ) were brought to room temperature or deparaffinized and washed in PBS. Antigen retrieval was performed in citrate buffer, pH 6.0 (Sigma-Aldrich) using a water bath (95°C for 10–12 min) or antigen retrieval system (Electron Microscopy Science). Sections were washed with 0.1% Triton in PBS (PBST), incubated in blocking buffer (1% BSA in 0.1% PBST) for 1 h, and then stained with primary antibody for 2 h at RT or overnight at 4°C. Following primary antibody incubation, tissues were washed three times in PBST followed by incubation with secondary antibody in blocking buffer for 1 h. Sections were washed with PBST, and coverslips were mounted with Fluor G reagent with DAPI. Images were captured with a confocal microscope using 20X or 40X objectives and visualized using 3D projection in Imaris software. For immunohistochemistry, tissue sections were deparaffinized/rehydrated, antigens were retrieved by citrate buffer, and non-specific binding was blocked. Sections were incubated with primary antibodies against perilipin, phosphorylated SMAD2, phospho-SMAD3, and PDGFβR, αSMA, CD31, CD206, and collagen.

### Real-time PCR

Total RNA was extracted from cells using RNeasy Mini Kit (QIAGEN). cDNA was prepared from the RNA (High-Capacity RNA-to-cDNA Kit; Applied Biosystems). Real-time PCR was performed with ready-made primer sets for mouse *collagen 1* and *ACTA2* genes using a real time PCR system (Bio-Rad). The thermal cycler conditions were 50°C for 2 minutes, followed by 95°C for 10 minutes and 95°C for 15 seconds, and finally 60°C for 1 minute. A total of 40 cycles were run. Data were normalized to the endogenous control gene GAPDH.

### Statistical analysis

Statistical analysis was performed using SAS version 9.4 (SAS Institute, Inc). We used the multiple linear regression model for fibrosis and diastolic dysfunction. For comparison, the Kruskal-Wallis test and pairwise 2-sample Wilcoxon rank sum tests were used. All data are expressed as mean ± standard deviation or standard error of the mean to determine whether differences were statistically significant. P < .05 was considered statistically significant.

## Acknowledgments

We thank P Dube and TR Dilling, W Ben for helping with ART administration, Immunoblotting, and tissue harvesting. We thank Drs. M. Zanni, T. Neilan, and A. Neilan for their comments editing of the manuscript. This work was fully supported by HL167656 (JA) and partly supported by HL148123 (JA) and the Angelo Donghia Foundation (JL).

## Notes

### Competing Interest Statement

The authors have declared no competing interest.

