## Supplemental Information for "Atorvastatin suppresses cardiac fibrosis and dysfunction induced by HIV and certain antiretroviral drugs in mice by blocking platelet TGFβ1"

[a]Cardiovascular Biology Research Program, Oklahoma Medical Research Foundation, Oklahoma City, OK 73104; [b]University of Oklahoma Health Sciences Center, Oklahoma City, OK 73104; [c]Division of Infectious Diseases, Weill Cornell Medical College, New York, NY 10065; [d]Baylor College of Medicine, Houston, TX 77030; and [e]Division of Hematology and Medical Oncology, Weill Cornell Medical College, New York, NY 10065.

**Running Title:** Role of platelet TGFβ1 in organ fibrosis

[1]To whom correspondence may be addressed. Jasimuddin Ahamed, Ph.D.

Fig. S1

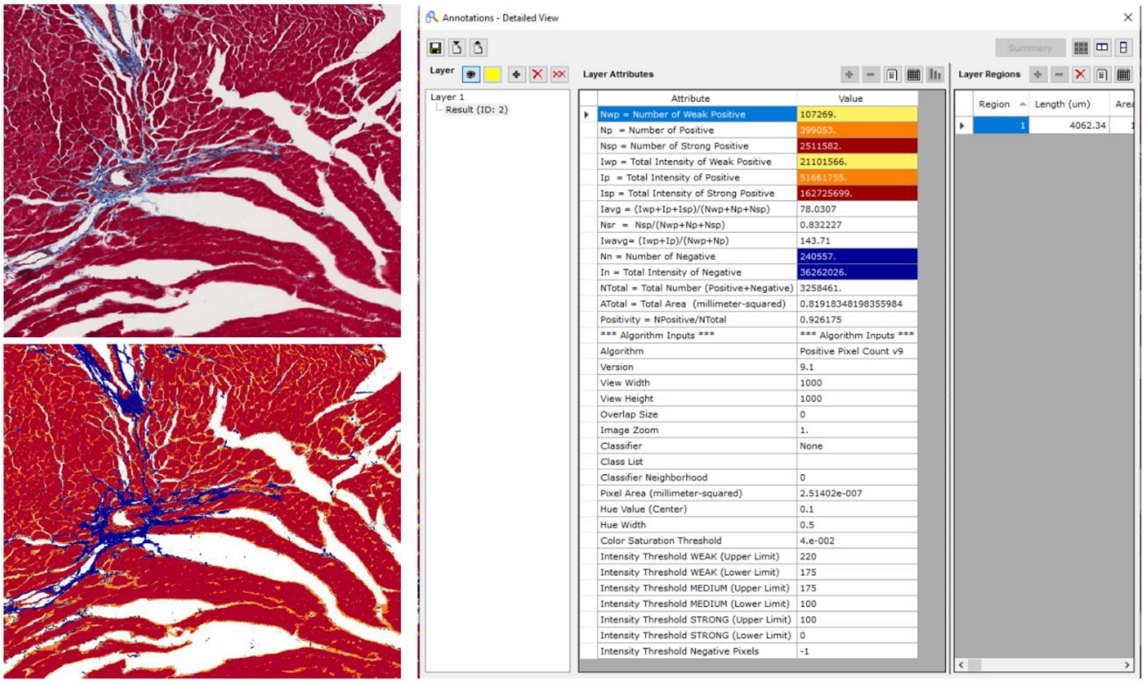

**Fig. S1.**

Masson trichome-stained collagen image showed blue color collagen areas with fibrosis (upper panel). For quantification of fibrosis, the whole heart section images were carefully examined to identify focal fibrosis and remove heart valves-associated with collagen staining, and lower panel image in the ImageJ program (NIH) showing adjustments were made and set an algorithm to calculate the percentage of fibrotic area within the total area of each heart section image.

Fig. S2

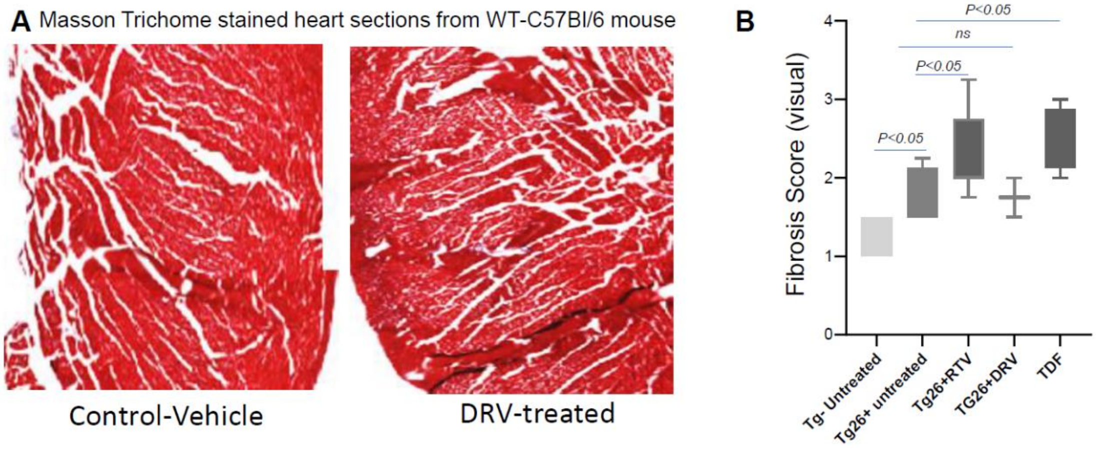

**Fig. S2.**

**(A)** Representative Masson Trichome images of vehicle-control and darunavir (DRV)-treated heart sections from C57Bl/6 mice. (**B**) Blind scoring of fibrosis by pathologists/tech, showing higher fibrosis in ART (RTV-, or TDF-FTC-DTD-, but not DRV-treated HIV-Tg26 compared to untreated control mice.

Fig. S3

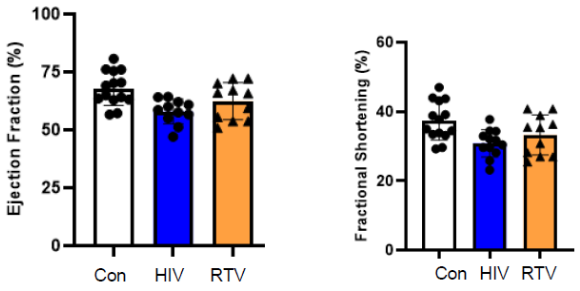

**Fig. S3.**
Systolic heart functions parameters, ejection fraction (EF) and fractional shortening (FS), in control and HIV-Tg26 mice treated with ART for 8 weeks, were calculated from parameters obtained by M-mode echocardiography imaging using Vevo 2100 ultrasound device analyzed by FUJIFILM-VisualSonics cardio software.

Fig. S4

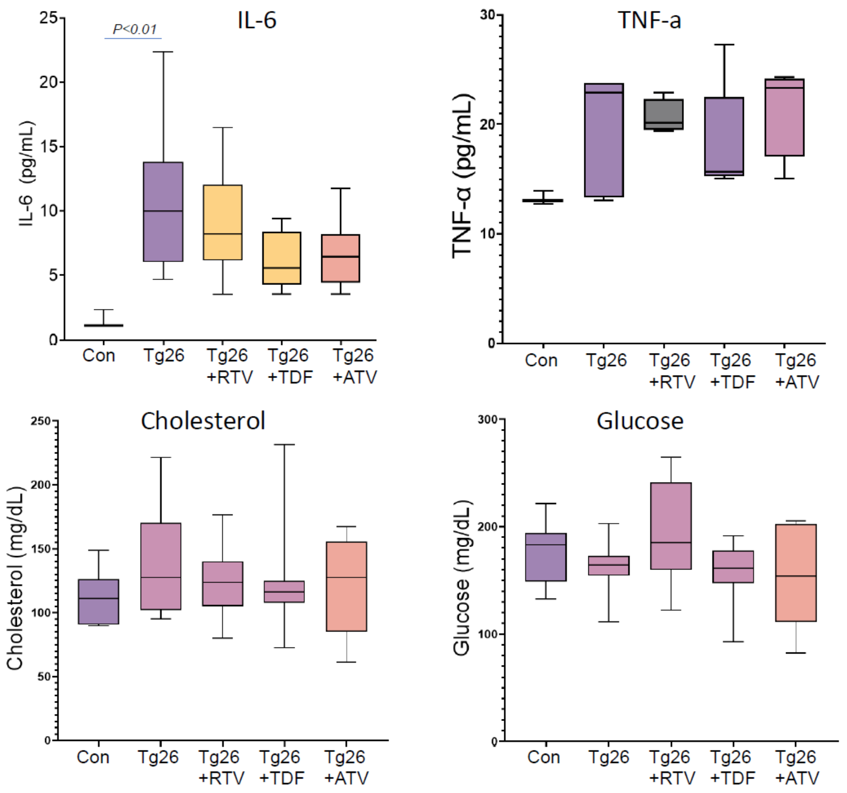

**Fig. S4.**

IL6 and TNFα in plasma from control (No-HIV) and HIV-Tg26 mice with and without ART (RTV or TDFc) and co-treated with ATV for 8 weeks, were measured by ELISA. Plasma total cholesterol and glucose levels were measured by colorimetric assays.

Fig. S5

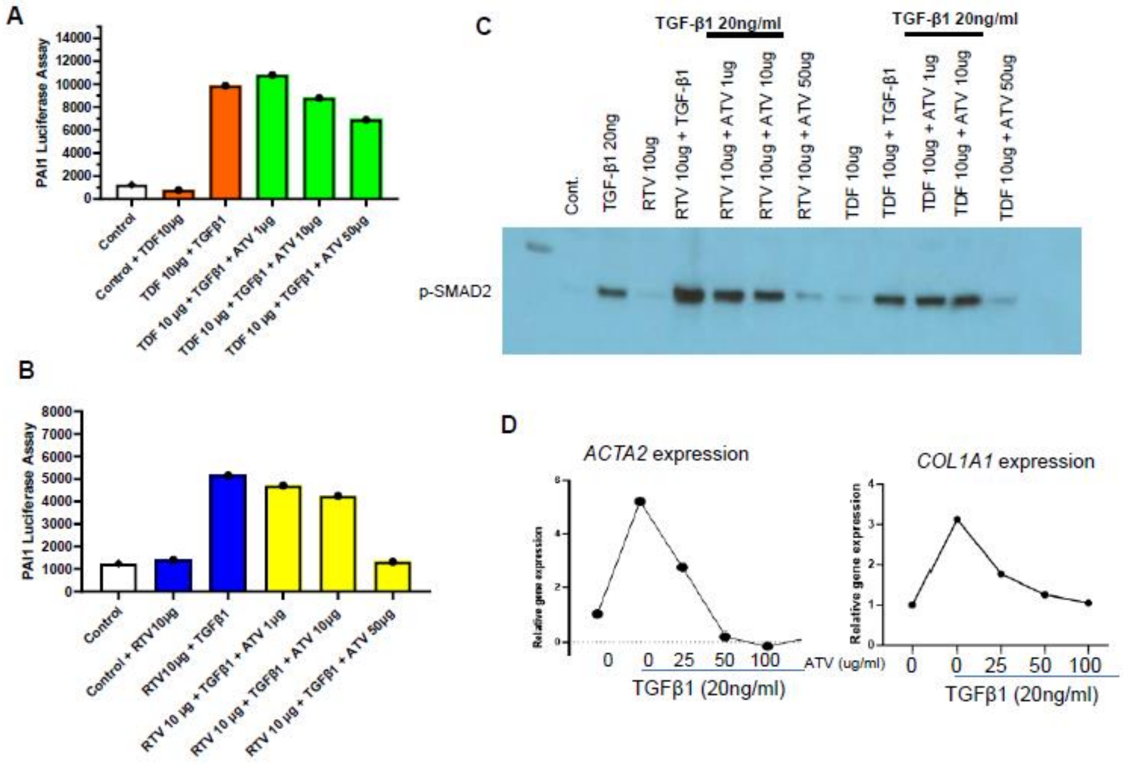

**Fig. S5.**

(**A-B**) Atorvastatin (ATV) inhibited TGFβ1 plus RTV- or TDF-mediated signaling for PAI-1 luciferase activity and (**C**) pSMAD2 phosphorylation in a dose-dependent manner. (**D**) *ACTA2* and *COL1A1* genes expression, which was inhibited by ATV as detected by real-time polymerase chain reaction stimulated with platelet-derived TGFβ1 for 3 hours.

Fig. S6

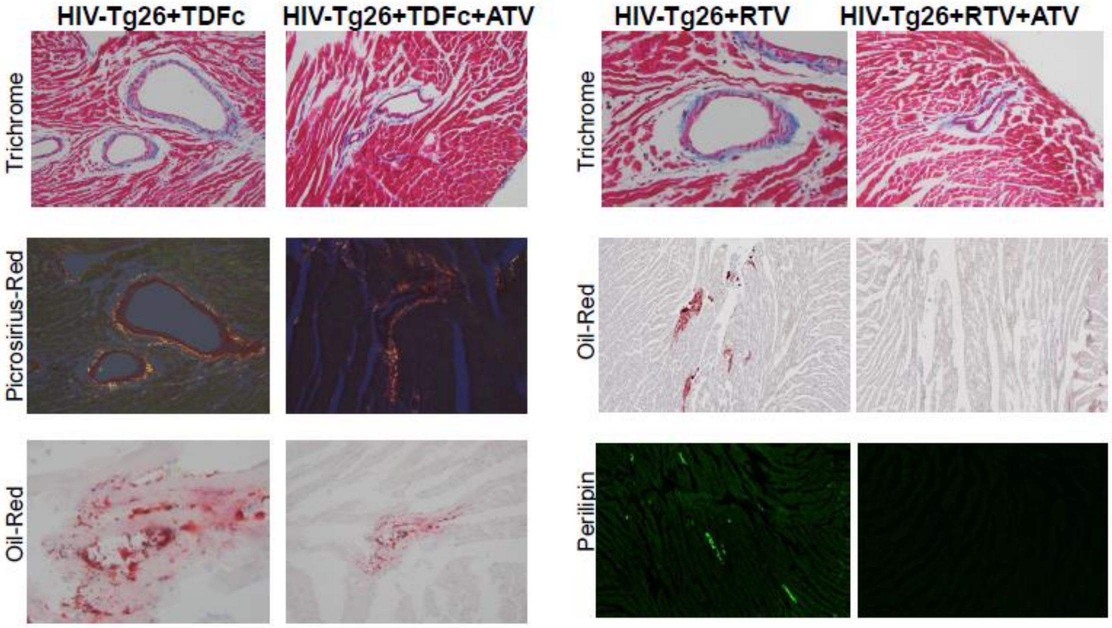

**Fig. S6.**

Atorvastatin (ATV) inhibited cardiac fibrosis, lipid/fat cells deposition as evidenced by Masson trichome, Picrosirius-Red, Oil-Red, and Perilipin staining in HIV-Tg26 mice.

### Fig. S7

Effect of ATV on ART-induced platelet activation and TGFβ1 release

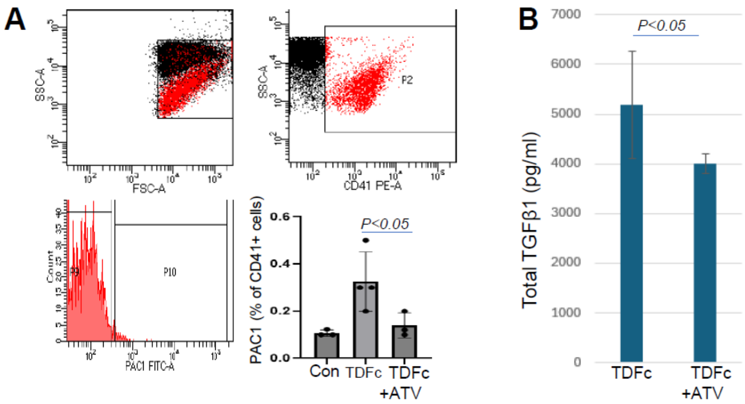

**Fig. S7.**
(**A**) Human platelet activation by TDFc was measured by flow cytometry gatting CD41-positive platelets expressing PAC1 binding before and after ATV treatment with TDFc. (**B**) Total TGFβ1 levels were measured in plasma from HIV-Tg26 mice exposed to TDFc and co-treated with ATV (TDFc+ATV).

Fig. S8

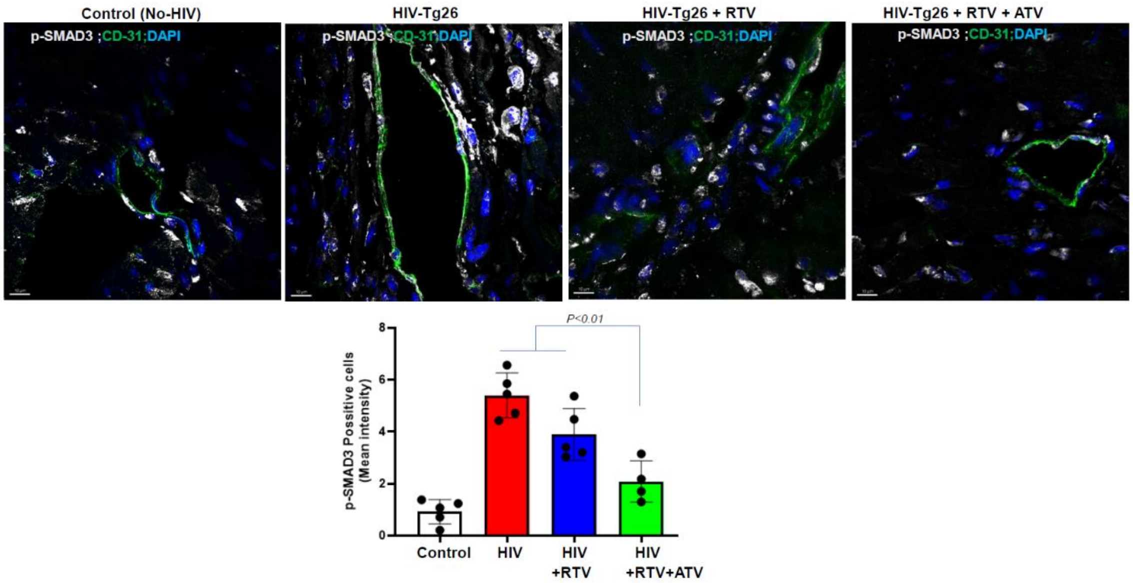

**Fig. S8.**

Atorvastatin (ATV) inhibits Smad signaling-representative confocal images [phosphorylation of pSMAD3 (white color) and pSmad2 (not shown)] and endothelial cells stained with CD31 antibody (green color) in heart of control (No-HIV), HIV-Tg26 mice treated with RTV with and without ATV. Lower panel shows the quantification of TGFβ signaling response assessed by pSmad3 staining intensity.

**<u>List of antibodies and reagents</u>**

| Antibodies/reagents | Vender | Cat No: | Dilution | Species | Note |
|---|---|---|---|---|---|
| Anti-pSMAD2 | Millipore | AB3849 | 1:1000 | Rabbit | Western blotting & Immunostaining |
| Anti SMAD2/3 | Cell Signaling | 8685 | 1:1000 | Rabbit | Western blotting |
| Anti-pSMAD3 | Rockland | 600-401-919S | 1:1000<br>1:250 | Rabbit | Western blotting & Immunostaining |
| Colllagen-1 | Sigma | SB1402151 | 1:1000 | Mouse | Immunostaining |
| Alpha -SMA | Thermofisher | 904601 | 1:250 | Mouse | Immunostaining |
| CD206 | Agilent | AF-2535 | 1:250 | Goat | Immunostaining |
| Perilipin-1 (K117) | Cell Signaling | 3467S | 1:250 | | Immunostaining |
| Donkey anti-Rabbit Alexa Fluor 647-conjugated | Jackson Immuno Research | 711-607-003 | 1:250 | Anti rabbit | Secondary antibody for IF |
| Donkey anti-goat Alexa Fluor 488-conjugated | Jackson Immuno Research | 705-546-147 | 1:250 | Anti goat | Secondary antibody for Immunostaining |
| Donkey anti-rat Alexa Fluor 594-conjugated | Jackson Immuno Research | 712-586-150 | 1:250 | Anti rat | Secondary antibody for Immunostaining |
| Donkey anti-mouse Alexa Fluor 594-conjugated | Jackson Immuno Research | 715-586-150 | 1:250 | Anti mouse | Secondary antibody for Immunostaining |
| Goat anti-Rabbit, HRP-conjugated | Jackson Immuno Research | 711-035-152 | 1:10000 | Anti rabbit | Secondary antibody for Western blotting |
| Protein ladder | Bio-Rad | 161-0374 | | | |
| TGF beta1 | Peprotech | 100-21-2ug | 20ng/mL | | |
| Oil Red stain kit | Newcomer Supply INC | 1277B | | | Paraffin section or Cryosection |
| Picrosirius Red Stain kit | IHC World | 3012 | | | Paraffin section or Cryosection |
| Masson Trichrome Stain kit | Electron Microscopy Science | 26367-01 | Bouin'S Fixative | | Paraffin section or Cryosection |
| | | 26367-02 | Weigerts Iron Hematoxylin A | | |
| | | 26367-03 | Weigerts Iron Hematoxylin B | | |
| | | 26367-04 | Biebrich Scarlet Solution 1%, | | |
| | | 26367-05 | Phosphomolybdic/Phosphotungstic Acid Soltn | | |
| | | 26367-06 | Aniline Blue Solution | | |
| | | 26367-06 | Acetic Acid 1% Aqueous | | |
| SYBR™ Select Master Mix | Applied Biosystems™ Thermofisher | 44-729-08 | RT-PCR | | |
| High-Capacity RNA-to-cDNA™ Kit | Applied Biosystems™ Thermofisher | 43-874-06 | RT-PCR | | |
| Primers for RT-PCR | IDT | Col1A1 F | 5-GAAACCCGAGGTATGCTTGA-3 | | |
| | | Col1A1 R | 5-GTTGGGACAGTCCAGTTCTT-3 | | |
| | | ACTA2 F | 5-CCTTCGTGACTACTGCCGAG-3 | | |
| | | ACTA2 R | 5-AATGCCTGGGTACATGGTGG-3 | | |
| | | GAPDH F | 5-GGCAAATTCAACGGCACAGT-3 | | |
| | | GAPDH R | 5-CGCTCCTGGAAGATGGTGAT-3 | | |
